# Delay-period activity in frontal, parietal, and occipital cortex tracks different attractor dynamics in visual working memory

**DOI:** 10.1101/2020.02.13.947887

**Authors:** Qing Yu, Matthew F. Panichello, Ying Cai, Bradley R. Postle, Timothy J. Buschman

**Affiliations:** Department of Psychiatry, University of Wisconsin–Madison, Madison, WI 53719, USA; Department of Psychology, University of Wisconsin–Madison, Madison, WI 53706, USA; Priceton Neuroscience Institute, Princeton University, Princeton, NJ 08540, USA; Department of Psychology, Princeton University, Princeton, NJ 08540, USA; Department of Psychology and Behavioral Sciences, Zhejiang University, Hangzhou, Zhejiang, 310007, China

## Abstract

One important neural hallmark of working memory is persistent elevated delay-period activity in frontal and parietal cortex. In human fMRI, delay-period BOLD activity in frontal and parietal cortex increases monotonically with memory load and asymptotes at an individual’s capacity. Previous work has demonstrated that frontal and parietal delay-period activity correlates with the decline in behavioral memory precision observed with increasing memory load. However, because memory precision can be influenced by a variety of factors, it remains unclear what cognitive processes underlie persistent activity in frontal and parietal cortex. Recent psychophysical work has shown that attractor dynamics bias memory representations toward a few stable representations and reduce the effects of internal noise. From this perspective, imprecision in memory results from both drift towards stable attractor states and random diffusion. Here we asked whether delay-period BOLD activity in frontal and parietal cortex might be explained, in part, by these attractor dynamics. We analyzed data from an existing experiment in which subjects performed delayed recall for line orientation, at different loads, during fMRI scanning. We modeled subjects’ behavior using a discrete attractor model, and calculated within-subject correlation between frontal and parietal delay-period activity and estimated sources of memory error (drift and diffusion). We found that although increases in frontal and parietal activity were associated with increases in both diffusion and drift, diffusion explained the most variance in frontal and parietal delay-period activity. In comparison, a subsequent whole-brain regression analysis showed that drift rather than diffusion explained the most variance in delay-period activity in lateral occipital cortex. These results provide a new interpretation for the function of frontal, parietal, and occipital delay-period activity in working memory.

## Introduction

Working memory – the ability to mentally retain and manipulate information to guide behavior – is crucial for many aspects of high-level cognition [1-3]. One prominent neural hallmark of working memory performance is persistent elevated delay-period activity in frontal and parietal cortex. Specifically, blood oxygen level-dependent (BOLD) activity in frontal and parietal cortex increases monotonically with memory load and asymptotes at an individual’s memory capacity [4, 5]. Activity in these networks is thought to reflect the engagement of control [6, 7]. For example, one recent study has demonstrated that persistent activity in parietal cortex tracks the demands of binding stimulus content to its trial-specific context, rather than memory load per se [8]. These signals have been shown to correlate with individual memory capacity [4, 5] and with memory precision [8-10]. In contrast, persistently elevated activity during the delay period is often absent in occipital cortex, despite the reliable representation of stimulus-specific information [8, 10-13].

Recent psychophysical work has shown that inaccuracies in working memory are due to both random error and systematic biases. For example, when subjects remember features drawn from a uniform stimulus space, their responses are not uniform. Instead, the responses “cluster” around a small number of specific values [14-16]. Further modeling work has demonstrated this clustering can be explained by attractor dynamics that pull memories to specific locations in mnemonic space (i.e. color memories are ‘attracted’ to red). While this induces systematic error into the memories, it also stabilizes memories near the attractors [16]. Thus, engaging attractor dynamics is thought to be especially beneficial when memory load is higher, because increased noise in stimulus representations can be counteracted by increasing drift towards a few stable representations.

Because load-related imprecision in working memory performance reflects both random diffusion and drift towards stable attractor states, it remains unclear which of these dynamics could account for load-sensitive delay-period activity in parietal and frontal cortex. In the current study, we analyzed data from an existing experiment in which subjects performed delayed recall for line orientation, at different memory loads, during fMRI scanning. We modeled subjects’ behavior using a discrete attractor model, and regressed the resultant load-sensitive estimates of drift and diffusion against load-dependent delay-period activity in parietal and frontal cortex. We found that an increase in frontal and parietal activity was associated with increases in both diffusion and drift. Furthermore, diffusion rather than drift explained the most variance in frontal and parietal delay-period activity. In comparison, a subsequent whole-brain regression analysis showed that drift rather than diffusion explained the most variance in delay-period activity in lateral occipital cortex. The results provided a novel interpretation of the functions associated with delay-period activity, suggesting frontoparietal control networks may be engaged to offset load-related diffusive noise while load-related drift is localized to occipital cortex.

## Results

### Behavioral performance

Subjects performed a delayed estimation task on line orientations. On different trials, subjects either remembered one orientation (*1O*), or three different orientations (*3O*). For subjects who participated in the fMRI sessions, we first plotted the distribution of their raw responses (*n* = 16), separately for *1O* and *3O* trials. Recall error, measured as the angular distance between the target orientation and response orientation, increased with increasing memory load, *t*(15) = 8.27, *p* = 5.68 × 10^−7^. Furthermore, similar to what has been previously reported for color [14-16], subjects’ responses to orientation working memory also clustered around a small number of orientations (Figure 1B).

**Figure 1A.**
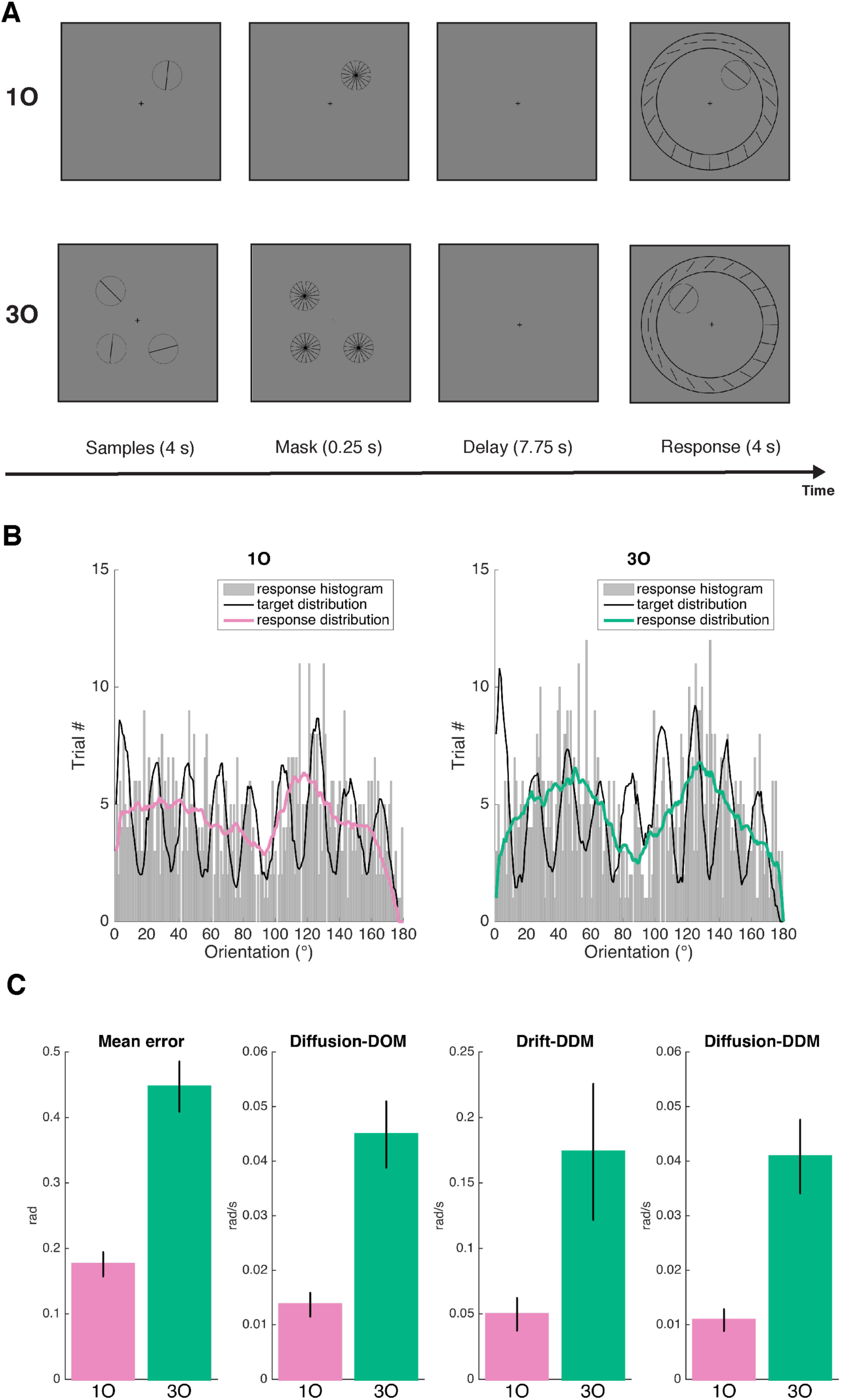
Trial sequence of the fMRI task and Behavioral performance. **A.** For the data analyzed in the current study, participants remembered either one orientation (*1O*), or three orientations (*3O*). Sample stimuli were presented on the screen for 4 s, followed by a brief mask period of 0.25 s. After a delay of 7.75 s, participants rotated the needle of the response wheel to indicate the remembered orientation at the probed location. **B.** The raw response distribution of *1O* and *3O* trials, indicated by the gray histograms. The black lines indicate the envelope of target distribution, and pink and green lines indicate the envelope of response distribution, for *1O* and *3O* trials separately. **C.** Model-free and model-based behavioral performance. From left to right panel shows mean error, diffusion from the DOM model, drift from the DDM model, and diffusion from the DDM model. Error bars indicate ± 1 SEM.

To account for these clusters, we fit the behavioral data with the drift-diffusion model (DDM), which included drift towards attractor locations. For comparison, we also fit the ‘diffusion-only’ model (DOM). Consistent with previous work on color working memory [16], the DDM provided a better fit to behavior than the DOM (difference in cross-validated log-likelihood = 3.67). For the DDM, the diffusion and the drift parameters both increased with memory load (*t*(15) = 4.86, *p* = 0.0002 and *t*(15) = 2.43, *p* = 0.028, respectively), as did the diffusion parameter from the DOM (*t*(15) = 6.52, *p* = 9.67 × 10^−6^; Figure 1C). When we repeated these analyses on the full set of behavioral data (*n* = 30; including behavior-only subjects), all results were qualitatively similar to those reported above (the average difference in cross-validated log-likelihood across folds was 6.56 between DDM and DOM).

### BOLD signal change in IPS and PFC

We next examined the BOLD time course in IPS and in PFC during the working memory task, at the two memory loads. We observed the classic pattern of load-sensitive BOLD activity in both ROIs: signal intensity was sustained above baseline across the delay period in both load conditions (all *p*s < 0.001), with greater activity for the higher memory load condition (all *p*s < 0.01, including the “late-delay” TR, at which BOLD-behavior analyses were carried out; Figure 2A and 2B).

**Figure 2.**
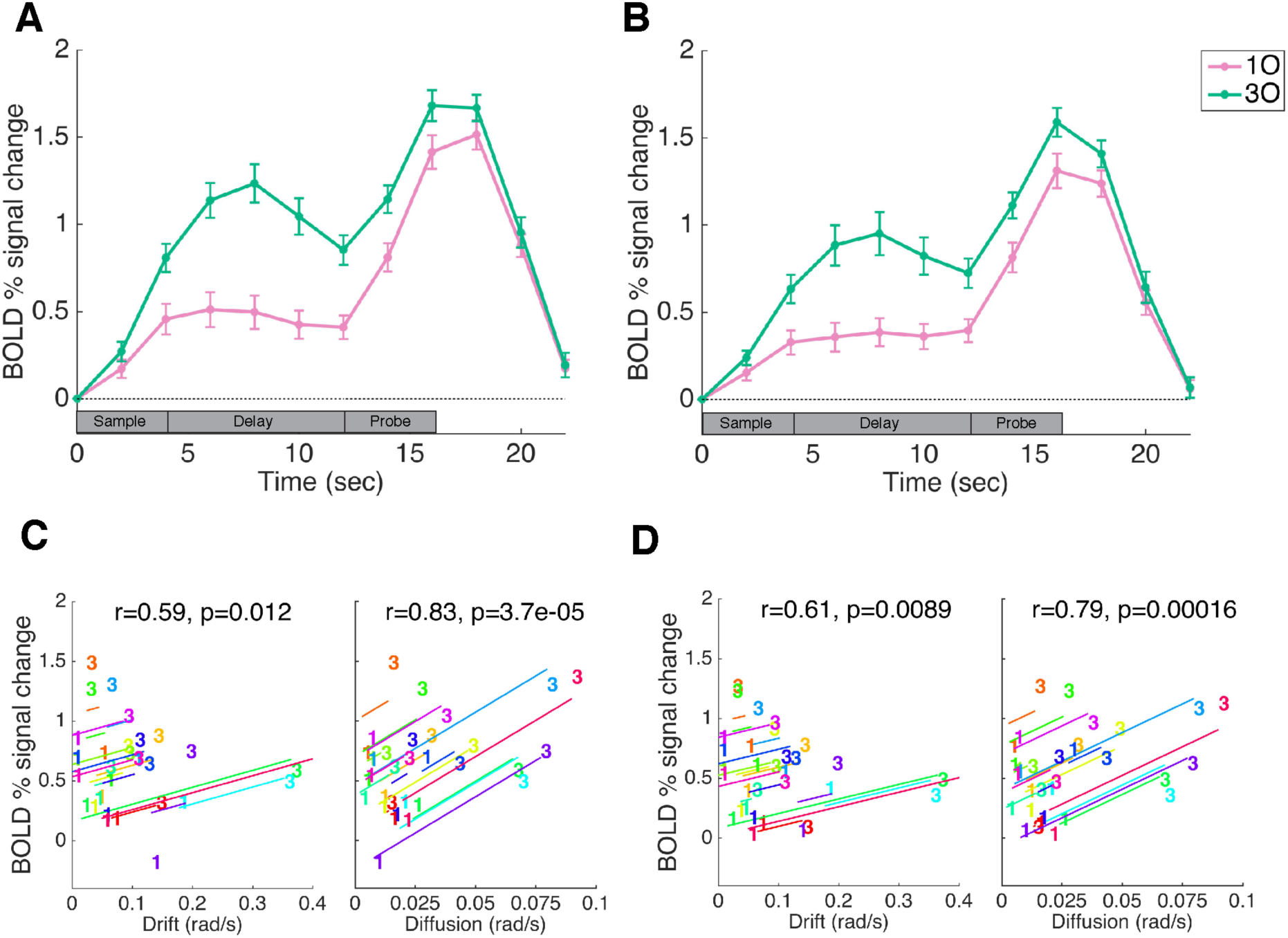
BOLD activity and brain-behavior correlations in IPS and PFC. **A**. Trial-averaged BOLD activity in the IPS functional ROI. **B**. Time course of BOLD activity in the PFC functional ROI. Pink and green lines correspond to the *1O* and *3O* conditions, respectively. Error bars indicate ± 1 SEM. **C**. Within-subject correlations between behavioral parameter from DDM (drift and diffusion plotted separately) and IPS BOLD activity, at “late delay” time point (12 s). **D**. within-subject correlations between behavioral parameter (drift or diffusion) and PFC BOLD activity. In each plot, data from each subject are plotted in a different color, and the “1” and “3” symbols correspond to values from *1O* and *3O* trials, respectively. Lines illustrate the best fit of the group-level linear trend (i.e., the within-subject correlation) in relation to individual subject data.

### Modeling load-dependent BOLD activity with behavior at the ROI level

To relate load-dependent BOLD activity in parietal and frontal cortex to behavior, we fitted linear regression models with behavioral-model fitted parameters and subject as the independent variables, and BOLD activity as the dependent variable. We first used these regression models to calculate within-subject correlations (ANCOVAs) between behavioral parameters (drift and diffusion) and BOLD activity. The results indicated that BOLD activity in both ROIs correlated significantly with diffusion (IPS diffusion: *r* = 0.83, *p* = 0.00004; PFC diffusion: *r* = 0.79, *p* = 0.0002) and drift (IPS drift: *r* = 0.59, *p* = 0.012; PFC drift: *r* = 0.61, *p* = 0.009; Figure 2C and 2D).

Next, to evaluate the contribution of drift and diffusion, we found the regression model that best explained BOLD activity in the two ROIs. Comparison between the four models of interest indicated that Model 2 (BOLD ∼ diffusion (DDM) + subject) explained the most variance in BOLD activity in both IPS and PFC ROIs, and showed the best model performance in terms of AIC and BIC (See Table 1 for a complete list of model comparisons).

**Table 1:** Comparison between different regression models.

| Model | adjusted $R^2$ | AIC | BIC |
| --- | --- | --- | --- |
| IPS |  |  |  |
| Model 1 | 0.237 | 29.3792 | 54.2967 |
| Model 2 | 0.635 | 5.7543 | 30.6718 |
| Model 3 | 0.619 | 6.9660 | 33.3493 |
| Model 4 | 0.580 | 10.2552 | 35.1757 |
| **PFC** |  |  |  |
| Model 1 | 0.412 | 14.1163 | 39.0338 |
| Model 2 | 0.659 | -3.2714 | 21.6461 |
| Model 3 | 0.652 | -2.8174 | 23.5658 |
| Model 4 | 0.566 | 4.3899 | 29.3075 |

We also used stepwise regression to examine the relative contribution of drift and diffusion to the prediction of BOLD activity. Starting from Model 3 (BOLD ∼ drift (DDM) + diffusion (DDM) + subject), stepwise regression removed drift from the model for both IPS (*F*(1,14) = 0.35, *p* = 0.564) and PFC (*F*(1,14) = 0.84, *p* = 0.376), but retained diffusion for both ROIs (diffusion vs. constant model: IPS: *F*(32,15) = 4.37, *p* = 0.003; PFC: *F*(32,15) = 4.36, *p* = 0.003). Together, these results suggest the level of BOLD activity in both IPS and PFC is most strongly correlated with the amount of diffusive noise in memories.

### Modeling load-dependent BOLD activity with behavior at the whole-brain level

Lastly, we performed a whole-brain linear regression analysis to explore the relative contribution of drift and diffusion to the BOLD activity of each voxel. Consistent with our ROI-based results, we found significant clusters in bilateral IPS and left frontal cortex with load-dependent BOLD activity that can be better explained by load-dependent changes in diffusion (Figure 3A, red clusters).

**Figure 3.**
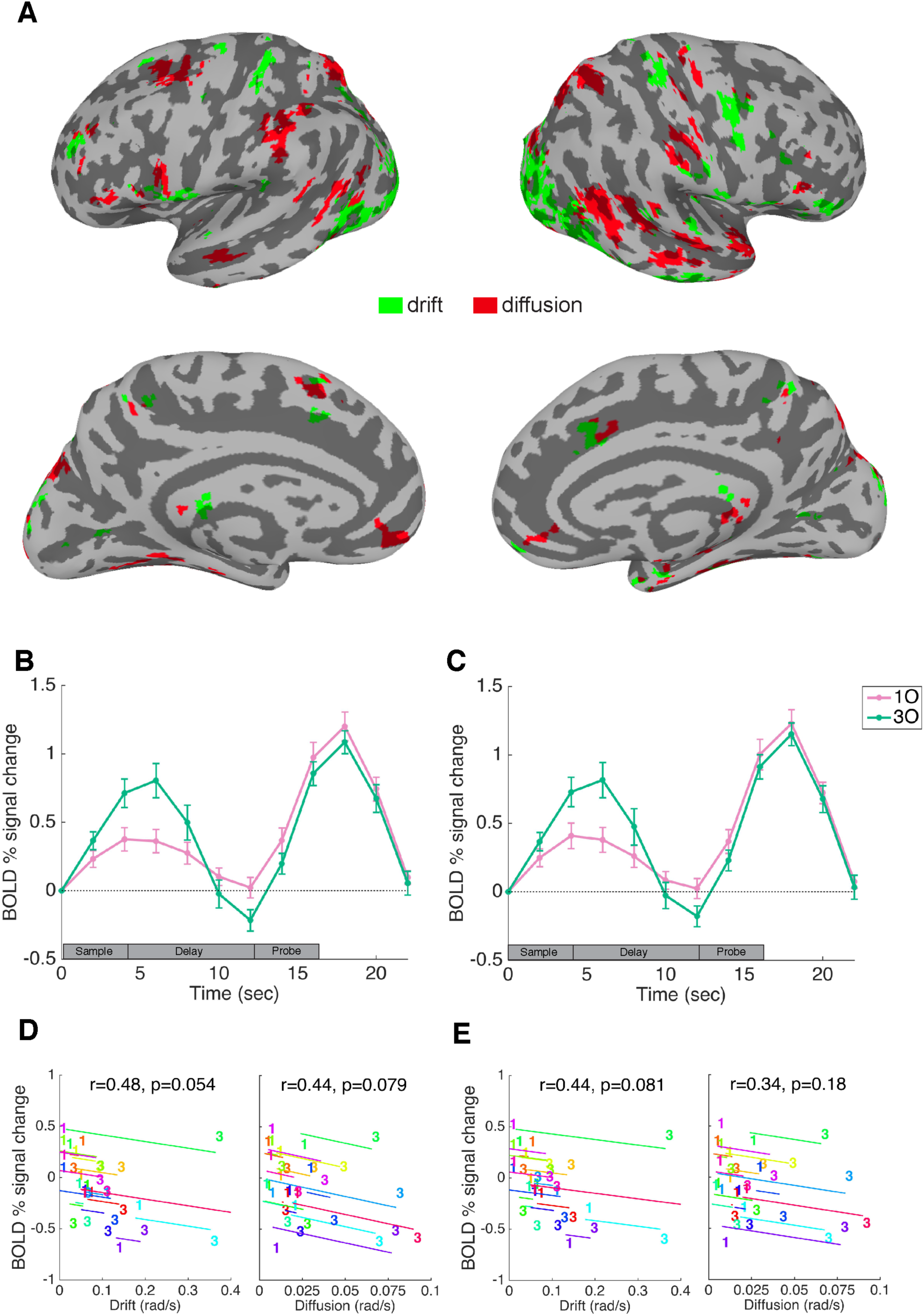
Whole-brain regression analysis with drift and diffusion and ROI-based results in LO. **A.**Whole-brain regression with drift and diffusion. Green denotes voxels showing load-dependent BOLD activity that can be better explained by load-dependent changes in drift, and Red denotes voxels showing load-dependent BOLD activity that can be better explained by load-dependent changes in diffusion. For visualization purposes, results were clusterized at a threshold of 20 voxels. The left two panels show results from the left hemisphere, and the right two panels show results from the right hemisphere. The significance of the regression models was corrected using the FDR method at *p* < 0.05. **B.**Trial-averaged BOLD activity in the LO1 anatomical ROI. **C**. Time course of BOLD activity in the LO2 anatomical ROI. Pink and green lines correspond to the *1O* and *3O* conditions, respectively. Error bars indicate ± 1 SEM. **D**. Within-subject correlations between behavioral parameter from DDM (drift and diffusion plotted separately) and LO1 BOLD activity, at “late delay” time point (12 s). **E**. within-subject correlations between behavioral parameter (drift or diffusion) and LO2 BOLD activity. In each plot, data from each subject are plotted in a different color, and the “1” and “3” symbols correspond to values from *1O* and *3O* trials, respectively. Lines illustrate the best fit of the group-level linear trend (i.e., the within-subject correlation) in relation to individual subject data.

Interestingly, we also observed clusters that showed higher brain-behavior correlation with drift (Figure 3A, green clusters). These clusters were most prominent in in the lateral occipital cortex (LO), in superior postcentral gyrus bilaterally and in right inferior precentral gyrus. Because of the known involvement of occipital cortex in visual working memory, we defined two anatomical ROIs for LO (LO1 and LO2) and repeated with them the ROI-based analyses as previously performed for IPS and PFC.

Consistent with previous findings [8, 10-13], BOLD signal intensity in the two LO ROIs returned to baseline during the delay period, with late-delay period activity no different from baseline on *1O* trials (LO1: *t*(15) = 0.300, *p* = 0.868; LO2: *t*(15) = 0.315, *p* = 0.845) and slightly below-baseline on *3O* trials (LO1: *t*(15) = 2.754, *p* = 0.021; LO2: *t*(15) = 2.369, *p* = 0.043; Figure 3B and 3C). ANCOVAs between the behavioral parameters from the DDM and this BOLD activity revealed trending correlations with drift (LO1: *r* = -0.48, *p* = 0.054; LO2: *r* = - 0.44, *p* = 0.081) and less so with diffusion (LO1: *r* = -0.44, *p* = 0.079; LO2: *r* = -0.34, *p* = 0.18; Figure 3D and 3E). Furthermore, stepwise regression on Model 3 removed diffusion from the model for both LO1 (*F*(1,14) = 0.59, *p* = 0.456) and LO2 (*F*(1,14) = 0.13, *p* = 0.727), while drift remained in models for both ROIs (drift vs. constant model: LO1: *F*(32,15) = 3.98, *p* = 0.005; LO2: *F*(32,15) = 4.2, *p* = 0.004). This result was opposite of what was observed in the IPS and PFC ROIs.

## Discussion

The results of this study provide a new account of the function of load-sensitive activity in IPS and PFC [4, 5]. First, consistent with previous work with color working memory, here we showed that attractor dynamics provided a better account of behavioral data of orientation working memory, compared with classic mixture models that did not take attractor biases into account. Next, and most importantly, the diffusion parameter from the discrete attractor model provided the best account of the load-sensitive delay-period activity of IPS and PFC. In contrast, in LO where aggregate levels of late delay-period activity were at or below baseline levels, load-sensitive fluctuation in this activity was better explained by drift. Thus, our results provide the first evidence to our knowledge that load-related imprecision in working memory, known to entail increases in random diffusion and in drift towards stable attractor state, engages control-related circuits of IPS and PFC and sensory-related circuits of LO, respectively.

By definition, working memory is guided by information specific to the current trial. Nevertheless, working memory is also often influenced by many other factors, such as sensory history [17] and prior knowledge. In working memory for color, the influence of prior knowledge is reflected as clustered responses around a small number of specific color values, even when the distribution of sample colors is uniform [14-16]. The present results show that this phenomenon generalizes to another low-level visual feature, orientation, and these biases increased with increasing memory load. Together with those of Panichello et al. (2019), our results indicate that dynamical systems offer a useful framework within which to understand the influence of trial-nonspecific factors on working memory performance.

Neurally, delay-period neural activity in IPS and PFC increased with increasing memory load, and we showed that this load-dependent change in BOLD activity was mainly related to load-dependent changes in diffusion rather than drift. Therefore, load-related activity change in IPS and PFC is likely related to random diffusion processes, rather than systematic biases towards attractors. The random noise could be related to noise in representations when memories are held in IPS/PFC or related to greater engagement of control processes when working memory has greater diffusion. For example, a recent study has found that delay-period activity in IPS is more sensitive to the demands of context binding than of memory load per se. By this account, increases in diffusion were likely due, at least in part, to increased interference between representations of stimulus content and stimulus context, which would be expected to place greater demands on a frontoparietal priority map controlling visually guided behavior [8]. In comparison, load-related activity in LO was more sensitive to load-related changes in drift to particular stimulus values, rather than diffusion. This result is consistent with the idea that prior knowledge shapes feature tuning in visual cortex, resulting in biased tuning responses to different visual features at early stages of cortical processing [18].

When considering these findings, it is important to not think of these factors as working in isolation. In frontoparietal cortex, for example, estimating drift is still necessary, as it allows for a more accurate model of diffusion, that can better predict neural signals in these regions. Moreover, it is important to note that in terms of parameter fitting, the drift parameter relies inferring the shape of attractor landscape across the entire stimulus space, and therefore both the number of trials and the uniformity of target distribution can have a significant impact on the fitted outcome. It is possible that future studies acquiring more trials, and/or applying more uniformly distributed targets, will lead to improved model fit of drift, and increases in the variance explained by this parameter.

In previous studies emphasizing stimulus-specific representations of visual working memory, we have argued that disparate patterns of results in frontoparietal versus occipital cortex are consistent with a functional distinction between these two regions, with the former more strongly associated with control and the latter with stimulus representation [8, 10]. Here, we see that stimulus-nonspecific factors, as reflected in the relationship between load-dependent changes in behavior (drift and diffusion) and delay-period activity, are also consistent with this distinction. Taken together, the results from higher-order frontal and parietal cortex and low-level occipital cortex suggest that imprecision in working memory can be caused by a combination of effects of noise in parietal and frontal cortex, and of stimulus-related biases in occipital cortex.

## Method

### Subjects

The results reported here are from analyses carried out on existing data collected for other purposes [19, 20]. Thirty individuals (10 males, mean age 20.7 ± 2.3 years) participated in the behavioral session of the study, and sixteen of these (8 males, mean age 20.6 ± 1.8 years) also participated in two subsequent fMRI scanning sessions. All were recruited from the University of Wisconsin–Madison community. All had normal or corrected-to-normal vision, reported no neurological or psychiatric disease, and provided written informed consent approved by the University of Wisconsin–Madison Health Sciences Institutional Review Board. Anatomical scans from the fMRI session were also screened by a neuroradiologist, and no abnormalities were detected. All subjects were monetarily compensated for their participation.

### Stimuli and procedure

All stimuli were created and presented using Matlab (MathWorks, Natick, MA) and Psychtoolbox 3 extensions [21, 22]. In the behavioral session, stimuli were presented at a viewing distance of 62 cm on an iMac screen, with a refresh rate of 60 Hz. Subjects registered behavioral responses on a trackball response pad. In the fMRI session, stimuli were projected onto a 60-Hz Avotec Silent Vision 6011 projector (Avotec, Stuart, FL), and viewed through a coil-mounted mirror in the MRI scanner at a viewing distance of 69 cm. Subjects registered behavioral responses on a MR-compatible trackball response pad (Current Designs Inc., Philadelphia, PA).

There were three types of stimuli: oriented bars, color patches, or luminance patches. Each oriented-bar stimulus appeared as a black line (width = 0.08°) bisecting a white circle (radius = 2°). Line orientations were drawn from a pool of 9 orientations ranging from 0 to 160°, in 20° increments, with a random jitter of 1-5° added to each stimulus. Color patches were circular patches (radius = 2°) filled with one color drawn from a pool of 9 colors that were equidistant in CIEL*a*b color space (L = 70, a = 20, b = 38, radius = 60°), with a random jitter of 1-5°. Luminance patches were rendered as a gray circular patch (radius = 0.83°) inside a white annulus (radius = 2°), and the luminance of the patches were drawn from 9 grayscale values from [0.03, 0.03, 0.03] to [0.97, 0.97, 0.97], in steps of 0.1175. Throughout the experiment, the background screen color was gray [0.5, 0.5, 0.5].

There were three different trial types. On “*1O*” trials, one oriented bar was presented at one of four possible locations (45°, 135°, 225°, 315° relative to central fixation, with an eccentricity of 5°) for 4 s. Stimulus offset was followed by a mask (white circle [radius = 2°] bisected by 18 black bars [width = 0.08°] intersecting at their midpoints and each differing in orientation from its neighbors by 10°; 0.25 s) and a delay period (7.75 s) during which subjects maintained central fixation. Recall was prompted by the onset of a stimulus circle appearing at the same location as the sample, a response wheel centered on fixation (inner radius = 7.2°, outer radius of 9.2°), and a cursor (a conventional “mouse” arrow) located at central fixation. Twenty oriented lines (radius = 1.8°, width = 0.05°, ranging in orientation from 0° to 171° in steps of 9°) were displayed with equal spacing along the response wheel, and subjects registered their memory of the sample orientation by moving the cursor to the appropriate location on the response wheel and registering that location with a button press. At the onset of the recall display, the stimulus patch was rendered with a randomly determined value rendered in the format of the sample stimuli, and as soon as the subject began to move the cursor (with the trackball) the stimulus patch took on the value corresponding to the location on the response wheel that was nearest to the cursor. Responses were required within 4 s, while the circle and wheel remained on the screen. The angle of rotation of the response wheel was randomized across trials, to prevent subjects from preparing their response during the delay period.

“*3O*” trials were similar to “*1O*” trials, except three oriented bars, each with a different orientation, were displayed in three of the four possible sample locations, and, at time 12 s, the sample to be recalled was indicated by the location of the stimulus circle in the recall array. For each *3O* trial, sample values were selected randomly, without replacement, from the pool of 9 possible orientations (Figure 1A).

On “*1O1C1L*” trials, 1 oriented bar, 1 color patch, and 1 luminance patch were presented, and during the response stage subjects were tested, unpredictably, on their memory for one of these stimuli. The response wheel for color and luminance was the same size as the orientation wheel, but displayed 180 possible color or luminance values.

The behavioral session contained two blocks of *1O* and *3O* trials, and three blocks of *1O1C1L* trials. Each block contained 50 trials, and block order was counterbalanced across subjects. The *1O* and *3O* blocks contained 25 trials each for *1O* and *3O*, and the *1O1C1L* blocks contained 17 probes of two of the three categories, and 16 of the remaining one. The selection of the categories was randomized across blocks, yielding 50 trials for each category across three blocks.

There were two fMRI scanning sessions. The first scanning session included four 18-trial blocks of 9 *3O* trials and 9 *1O1C1L* trials (with 3 probes each for orientation, color, and luminance), yielding a total of 36 trials for each of these load-of-3 trial types. These four blocks were followed by eight 18-trial blocks of *1O* trials. The second session included twelve blocks of *1O* trials. To match the number of trials between conditions in fMRI data, two of the twenty *1O* blocks were randomly selected for each subject for further analyses.

We introduce the *1O1C1L* condition here only for the completeness of experimental design. All subsequent analyses focused on *1O* and *3O* trials for load-related changes in behavioral and neural data.

### Behavioral modeling

We fitted data from the behavioral session using a discrete attractor model [16]. This circular drift-diffusion model (DDM) fits the dynamic evolution of memories with two distinct processes: random noise (*diffusion*); and systematic *drift* towards one of several stable attractors. Notably, when the drift parameter is removed, the remaining diffusion-only model (DOM) is equivalent to a classic mixture model [23]. Both parameters are rates, with a unit of rad/s indicating the rate of diffusion and the maximum instantaneous drift rate. Unlike the Panichello et al. (2019) study, here we fitted behavioral data without separating out encoding and delay processes, because the length of memory delays was not manipulated in this experiment. The comparison between performance of the DDM and DOM models was evaluated by computing a 10-fold cross-validated log-likelihood value.

### fMRI Data acquisition

Whole-brain images were acquired with a 3 Tesla GE MR scanner (Discovery MR750; GE Healthcare, Chicago, IL) at the Lane Neuroimaging Laboratory at the University of Wisconsin–Madison HealthEmotions Research Institute (Department of Psychiatry). Functional images were acquired with a gradient-echo echo-planar sequence (2 sec repetition time (TR), 25 msec echo time (TE), 60° flip angle) within a 64 × 64 matrix (40 sagittal slices, 3.5mm isotropic). Each of the fMRI scans generated 215 volumes. A high-resolution T1 image was also acquired for each session with a fast spoiled gradient-recalled-echo sequence (8.2 msec TR, 3.2 msec TE, 12° flip angle, 172 axial slices, 256 × 256 in-plane, 1.0 mm isotropic).

### fMRI Data preprocessing

Functional MRI data were preprocessed using AFNI (http://afni.nimh.nih.gov) [24]. The data were first registered to the first volume of the first run, and then to the T1 volume of the first scan session. Six nuisance regressors were included in GLMs to account for head motion artifacts in six different directions. The data were then motion corrected, detrended (linear, quadratic, cubic), converted to percent signal change, and spatially smoothed with a 4-mm FWHM Gaussian kernel. For the whole-brain analysis, the data were further aligned to the MNI-ICBM 152 space [25].

### Region of interest (ROI) definition

We first defined anatomical ROIs using existing anatomical atlases, and warped them back to each subject’s structural scan in native space. Parietal anatomical ROIs were created by extracting intraparietal sulcus (IPS) masks IPS0-5 from the probabilistic atlas of Wang and colleagues [26], merging them, and collapsing over the right and left hemispheres. Lateral prefrontal cortex (PFC) anatomical ROIs were created by extracting masks of the superior, middle, and inferior frontal gyri supplied by AFNI, merging them, and collapsing over the right and left hemispheres. Lateral occipital anatomical ROIs were created by extracting masks for LO1 and LO2, from the probabilistic atlas of Wang and colleagues [26], merging them, and collapsing over the right and left hemispheres.

To find the functionally activated voxels within the anatomical atlases, a conventional mass-univariate general linear model (GLM) analysis was implemented in AFNI, with sample, delay and probe periods of the task modeled with boxcars (4 sec, 8 sec, and 4 sec in length, respectively) that were convolved with a canonical hemodynamic response function. Across the whole brain, we identified the 2000 voxels displaying the strongest loading on the contrast [delay -baseline], collapsing over all three conditions. The intersection of these 2000 voxels and the two anatomical masks defined the two functional ROIs in subsequent analyses: the IPS ROI and the PFC ROI. On average, the IPS functional ROI contained 463 ± 177 voxels, the PFC functional ROI contained 314 ± 86 voxels; the two anatomical LO ROIs contained 404 ± 57 and 456 ± 69 voxels, respectively.

### Univariate analyses

We calculated the percent signal change in BOLD activity relative to baseline for each time point during the working memory task; baseline was chosen as the average BOLD activity of the first TR of each trial. The BOLD signal change was averaged across trials within each condition, and across all voxels within each ROI. Statistical significance of BOLD activity against baseline was assessed using two-tailed, one-sample t-tests against 0, and the obtained *p* values were corrected across loads and time points using FDR (False Discovery Rate) [27]. Statistical difference of BOLD activity between *1O* and *3O* at each time point was assessed using two-tailed paired t-tests, and similarly the obtained *p* values were FDR corrected across time points.

### Brain-behavior correlation and model comparisons

Following previous work [8-10], we used an analysis of covariance (ANCOVA) method to evaluate the correlated sensitivity to trial type (i.e., *1O* vs. *3O*) across pairs of task-related variables (i.e., BOLD activity vs. behavioral parameter). Unlike simple correlations, ANCOVA accommodates the fact that each subject contributes a value for each level of trial type. It removes between-subject differences and assesses evidence for “within-subject correlation” between the two task-related variables [28].

Mathematically, within-subject correlations were implemented as linear regression models, and were calculated for *drift* and *diffusion* separately, where *subject* is a dummy variable for trial types (*1O* and *3O*) of each subject, and *BOLD* is BOLD signal from time 12 s (“late delay-period” activity):

Model 1: BOLD ∼ drift (DDM) + subject;

Model 2: BOLD ∼ diffusion (DDM) + subject;

The within-subject correlation *r* for drift or diffusion was calculated as:

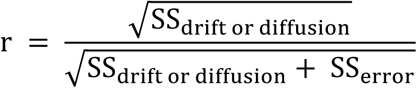

where SS stands for sum of squares.

To compare between the performance of different regression models, we included two more models, one full model that took both drift and diffusion into account, and one control model that used diffusion from the DOM model:

Model 3: BOLD ∼ drift (DDM) + diffusion (DDM) + subject;

Model 4: BOLD ∼ diffusion (DOM) + subject.

Model performance was evaluated by comparing Akaike Information Criterion (AIC), Bayesian Information Criterion (BIC), and adjusted *R*^2^ (explained variance of the model after adjusting for the number of predictors) of each model.

Lastly, we performed stepwise regression to evaluate the contribution of the drift and diffusion parameters to the prediction of BOLD activity. The regression model started with Model 3, after the initial fit, the predictors in the model were examined one by one, and the predictor with a *p* > 0.10 in the *F* test after removal was removed.

### Whole-brain regression analysis

To explore brain areas that showed activity sensitive to either the drift or diffusion parameter, we used a whole-brain exploratory analysis to find voxels with activity that can be best explained by either drift or diffusion. To this end, all subjects’ data were first normalized to the MNI-ICBM 152 space [25], and for each voxel we fit Models 1 and 2 to the BOLD activity of that voxel. The model with a higher adjusted *R*^2^ for each voxel was selected as the best fitting for that voxel, and we used the *p*-value of the selected model (*F*-test on regression vs. constant model) for statistical significance. To correct for multiple comparisons, we applied the False Discovery Rate (FDR) method to the *p*-values of the selected model across voxels. To avoid overinterpretation, we also applied a threshold in model selection using BIC [29], such that only voxels with a significant *p*-value after correction, and in which the drift or diffusion model outperformed the other by a BIC >= 2, remained in the final report. Therefore, we identified voxels with load-dependent BOLD activity that could be better explained by load-dependent changes in drift, or in diffusion, at the whole-brain level. Results from the whole-brain analysis were displayed on the cortical surface reconstructed with FreeSurfer (http://surfer.nmr.mgh.harvard.edu; [30, 31]) and visualized with SUMA in AFNI (http://afni.nimh.nih.gov) [24].

## Acknowledgements

This work was supported by NIMH R01MH064498 to B.R.P., and ONR N000141410681 and NIMH R01MH115042 to T.J.B.

